# Integrated meta-analysis of public human pancreatic single-cell transcriptomes reveals distinct beta-cell state trajectories across aging and diabetes

**DOI:** 10.64898/2026.01.27.701674

**Authors:** Elhadi Iich

## Abstract

Human pancreas single-cell RNA sequencing (scRNA-seq) studies have revealed extensive islet heterogeneity, yet cross-study integration remains limited by cohort- and platform-specific effects. Here, we assembled a unified atlas of >266,000 human pancreatic cells by harmonizing 18 publicly available scRNA-seq datasets spanning diverse technologies and donor phenotypes. Trajectory-based analyses resolved three beta-cell state trajectories associated with distinct stress axes. One trajectory reflects aging-associated transcriptional drift with progressive ER stress activation. A second captures diabetes-associated remodeling characterized by combined ER stress and induction of exocrine-like gene programs. A third highlights a metabolic-stress-associated program linked to lipid metabolism and polyhormonal transcriptional signatures in non-diabetic donors with elevated metabolic burden. In contrast to the relative stability of alpha-cell states, beta-cell identity programs eroded along specific trajectories, often preceding marked reductions in INS expression. Together, this integrated resource provides a scalable framework for dissecting human beta-cell plasticity and dysfunction using public single-cell transcriptomic data.

**HIGHLIGHTS:**

- Integrated atlas of >266,000 human pancreatic cells from 18 public scRNA-seq datasets spanning aging and diabetes
- Three beta-cell state trajectories associated with aging-related stress, diabetes-linked stress with exocrine-like program induction, and lipid-associated polyhormonal dedifferentiation
- Alpha-cell states are comparatively stable, whereas beta-cell identity programs erode along trajectory-specific paths, often preceding INS loss
- Diabetes-associated beta-cell subsets exhibit endocrine-exocrine transcriptional plasticity inferred from transcriptomic programs
- JUND identified as a candidate transcription factor associated with stress-linked exocrine gene expression in T2D beta cells

## INTRODUCTION

Understanding how human beta-cell dysfunction emerges across aging and type 2 diabetes (T2D) requires resolving the diversity of endocrine cell states in vivo. Single-cell RNA sequencing (scRNA-seq) has enabled cell-type-resolved profiling of transcriptional programs governing stress responses, identity maintenance, and functional plasticity within pancreatic islets^1,2^. However, deriving a unified biological framework from human islet scRNA-seq remains challenging, as observed heterogeneity reflects both genuine inter-individual variation and substantial technical differences across studies^3^.

Published human islet scRNA-seq datasets span diverse experimental designs, including plate-based and droplet-based platforms, variable tissue processing protocols, and distinct computational pipelines. These differences systematically affect transcript detection, stress-response signatures, and apparent substructure within endocrine populations, complicating direct cross-study comparisons^4,5,6,7,8^. Consequently, biological patterns observed within individual cohorts may be confounded by platform-specific biases^8,9^.

Beyond technical variability, beta-cell states themselves exhibit considerable biological diversity. Prior studies have reported beta-cell programs associated with insulin secretion capacity, ER stress, unfolded protein response activation, proliferation, and other adaptive processes4,^10,11,12^. Pseudotime analyses suggest that many of these states exist along continuous trajectories rather than as discrete cell types, consistent with dynamic remodeling in response to physiological and metabolic demands^11^.

Chronic metabolic stress in T2D drives beta-cell dysfunction, but the dominant molecular routes toward failure in humans remain unresolved. Reported disease-associated changes include activation of stress pathways, erosion of beta-cell identity programs, and partial engagement of non-endocrine transcriptional programs^13,14,15,16,17,18^. Whether these features represent a single convergent path or multiple trajectories linked to distinct stress axes remains unclear.

Recent advances in computational integration now allow harmonization of scRNA-seq datasets across cohorts and platforms, enabling higher-powered identification of conserved cell states and trajectories^19–27^. However, a comprehensive, pancreas-wide integrated atlas that resolves human beta-cell state trajectories across aging and diabetes and connects them to underlying regulatory programs is still lacking.

Here, we integrate 18 publicly available human pancreatic scRNA-seq datasets into a unified atlas spanning diverse donor phenotypes and technologies. Using trajectory-aware analyses, we identify three beta-cell state trajectories associated with distinct stress axes: aging-associated stress, diabetes-linked stress with exocrine-like program induction, and metabolic stress linked to lipid-associated polyhormonal dedifferentiation. This framework provides a systems-level view of human beta-cell plasticity and nominates candidate regulatory programs underlying trajectory-specific dysfunction.

## RESULTS

### Integrated analysis of 18 human pancreatic scRNA-seq studies resolves canonical pancreatic cell types

To derive a unified view of pancreatic cell-state organization across studies, we integrated 18 publicly available human pancreatic scRNA-seq datasets encompassing 176 samples generated across diverse platforms, donor demographics, and disease states, including non-diabetic (ND), type 2 diabetes (T2D), autoantibody-positive (AAB), and type 1 diabetes (T1D) conditions^1,4,5,11,12,28–39^ (Figure S1A; Tables S1 and S2).

Marked heterogeneity in sequencing depth, detected gene counts, and mitochondrial read fractions was observed across studies. Quality control was therefore performed on a per-dataset basis using standard metrics to exclude low-quality cells and technical outliers (Figure S1B; Table S3). To further reduce technical artifacts, we applied two complementary doublet detection approaches (DoubletFinder and scDblFinder), which identified partially overlapping sets of putative doublets (Figure S1C). Cells flagged by either method were excluded from downstream analyses.

After integration, the combined dataset comprised 266,614 single cells and resolved into 12 transcriptionally distinct clusters (Figure S1D). One cluster exhibited atypically low transcriptomic complexity despite high transcript counts, consistent with residual technical artifacts (Figure S1E), and was excluded from further analysis. The remaining 11 clusters were each supported by cells from at least 15 independent studies (Figures S1F-S1H), with no systematic association between cluster identity and sequencing platform (Figures S1I and S1J), indicating effective correction of platform-specific batch effects.

Clusters were not segregated by disease status alone, as neither ND nor diabetic samples formed exclusive cell-type partitions (Figures S1K and S1L). However, exocrine-enriched clusters showed a higher contribution from T1D donors, consistent with endocrine cell loss in this condition. Differential gene expression analysis identified canonical markers corresponding to major pancreatic cell types (Figure S2A; Table S4). Exocrine clusters were defined by ductal (e.g., *KRT18, MMP7, SPP1*) and acinar (e.g., *PRSS2, REG1A/B*) programs, while endocrine clusters expressed established markers of alpha (*GCG*), beta (*INS, IAPP*), delta (*SST*), and pancreatic polypeptide (*PPY*) cells. Additional clusters corresponded to stellate, endothelial, immune, stress-associated, and cycling populations based on established marker expression.

To complement marker-based annotation, we trained a supervised scPred classifier using well-annotated reference datasets and applied it post-integration to avoid circularity. The classifier achieved high performance for major endocrine and exocrine cell types (AUC > 0.99; Figure S2B). Predictions for rare populations were interpreted conservatively and retained only when concordant with independent marker expression. Final cell-type assignments were determined by agreement between cluster-level marker enrichment and classifier predictions (Figure S2C).

Low-level expression of epsilon-cell markers (e.g., *GHRL*) was detected in small subsets of cells distributed across multiple clusters, consistent with either transcriptional plasticity or sparse representation within broader endocrine states. Based on marker enrichment and embedding position, these cells were annotated as epsilon-like within mixed clusters (Figures S2D and 1A-1B). Two additional clusters were annotated as bihormonal endocrine cells and cycling alpha cells based on co-expression of endocrine markers and enrichment of cell-cycle-associated transcripts.

Together, these results demonstrate that large-scale integration of public human pancreatic scRNA-seq datasets robustly recapitulates canonical endocrine, exocrine, and stromal populations, providing a stable foundation for downstream trajectory and regulatory analyses.

### Trajectory analysis resolves structured transcriptomic states in beta cells but not alpha cells

To assess continuous transcriptional variation associated with metabolic disease, we focused on beta and alpha cells from non-diabetic (ND) and type 2 diabetes (T2D) donors. Cells were subset from the integrated atlas (beta: 26,740 cells; alpha: 57,836 cells) and re-integrated using SCTransform, excluding datasets contributing fewer than 100 cells to ensure stable inference.

Trajectory-oriented clustering using Monocle3 identified seven transcriptionally distinct beta-cell clusters (Figure 2A) but only four alpha-cell clusters (Figure S3A). Beta-cell clusters showed clear stratification by donor age and BMI, with cluster 3 enriched for younger, lean donors, while remaining clusters spanned higher ages and BMIs (Figure 2B). In contrast, alpha-cell clusters exhibited minimal stratification by age, BMI, or study representation (Figures S3B and S3C), consistent with relative transcriptional stability.

**Figure 1:**
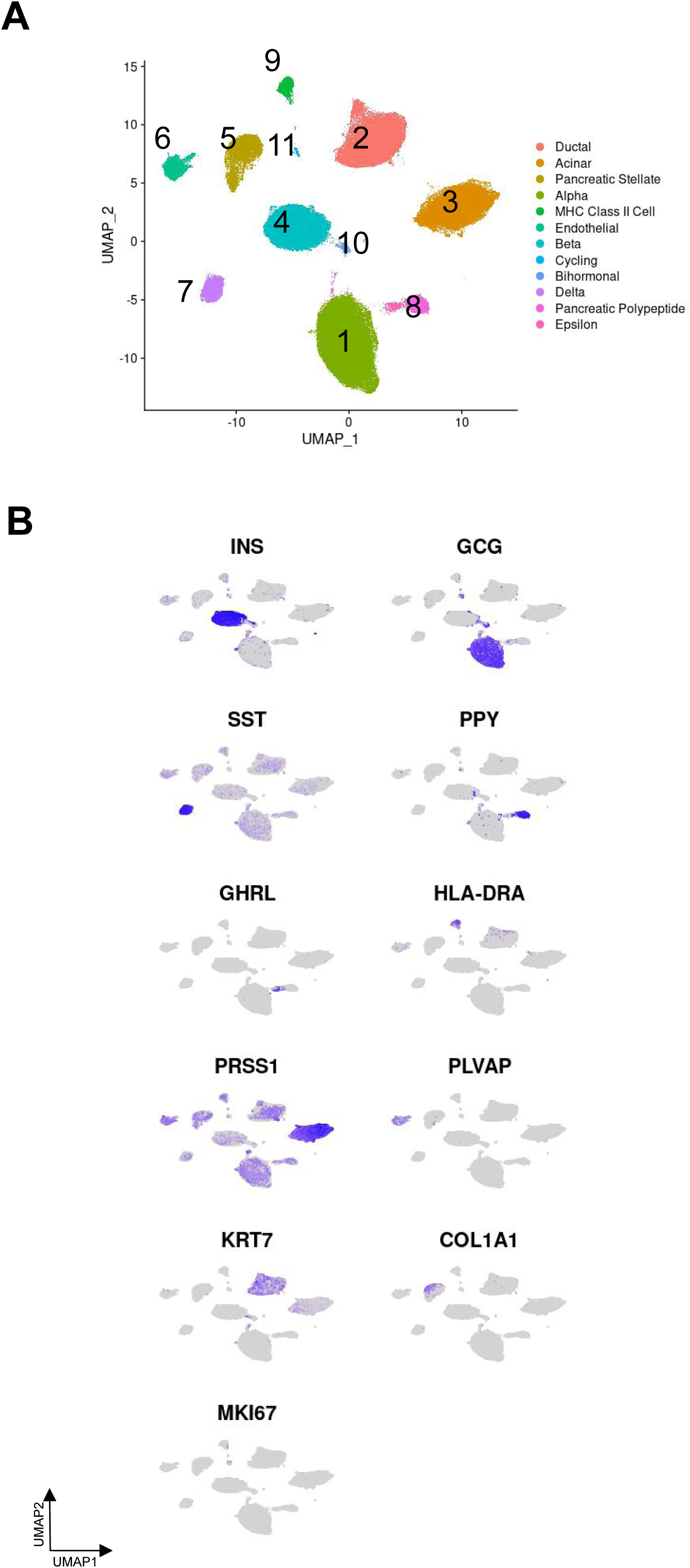
Integrated human pancreatic scRNA-seq atlas identifies major pancreatic cell subtypes. (A) Harmony integrated and scPred annotated scRNA-seq atlas. (B) Cell type marker expression of Insulin (*INS*), Glucagon (*GCG*), Somatostatin (*SST*), Pancreatic Polypeptide (*PPY*), Ghrelin (*GHRL*), HLA class II histocompatibility antigen DR alpha chain (*HLA-DRA*), Serine Protease 1 (*PRSS1*), plasmalemma vesicle associated protein (*PLVAP*), Keratin 7 (*KRT7*), Type 1 Collagen (*COL1A1*), and proliferation marker Ki-67 (*MKI67*).

**Figure 2:**
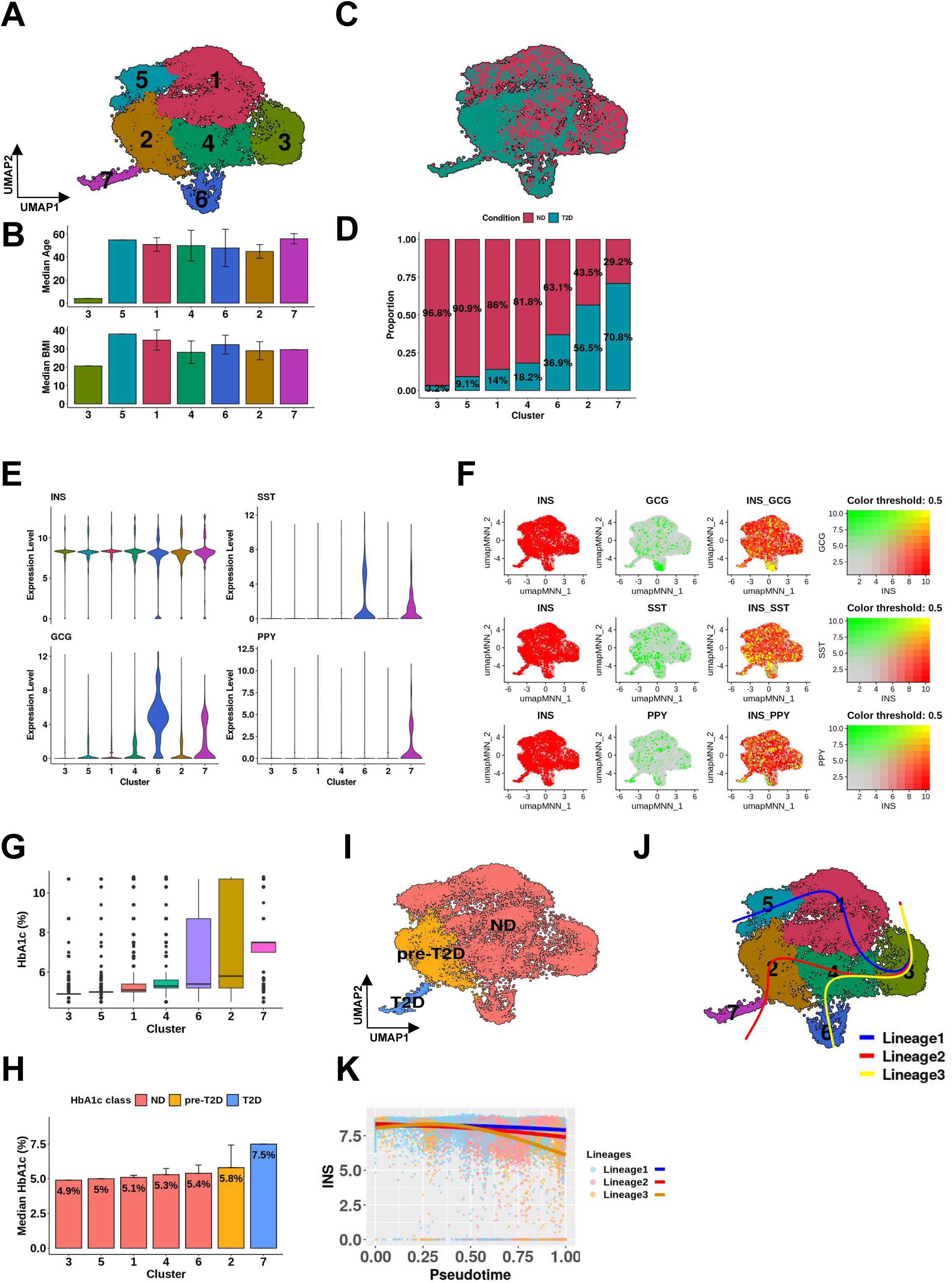
Integrated ND and T2D human beta-cells diverge into three different lineages. (A) UMAP representation of SCTransform integrated beta-cells with monocle3 identified clusters. (B) Median age and BMI of the beta-cell clusters. Error bars represent the Median Absolute Deviation (MAD). (C) Distribution and (D) proportion quantification of ND and T2D cells contributing to each beta-cell cluster. (E) Violin plots of *INS, GCG, SST* and *PPY* expression in each beta-cell cluster. (F) Visualization of co-expression of sets of markers *INS, GCG, SST* and *PPY* in beta-cells (yellow dots indicate co-expressing cells). (G) Distribution of HbA1c percentage across beta-cell clusters. (H and I) Beta-cell cluster subclassification based on median HbA1c. Error bars represent the MAD. (J) Slingshot inferred lineage trajectories starting from cluster 3 (predominantly ND cells with high *INS* expression). (K) Combined scatter plot and smoothers of *INS* expression changes along pseudotime for each beta-cell lineage. LOESS regression lines show smoothed expression trends.

Disease association analysis revealed a graded enrichment of T2D cells across beta-cell clusters, ranging from 3.2% to 70.8% (Figures 2C and 2D), defining a continuum from ND-to T2D-associated states. One ND-enriched cluster (cluster 6) exhibited co-expression of multiple endocrine hormones (*INS, GCG, SST*), consistent with previously described polyhormonal beta-cell states in non-diabetic donors (Figures 2E and 2F). By contrast, alpha-cell clusters showed limited disease stratification, with largely uniform *GCG* expression and only modest *INS* expression in a predominantly ND cluster (Figures S3D and S3E).

Incorporation of donor HbA1c values enabled classification of beta-cell clusters into ND, pre-T2D, and T2D states. Cluster 2 exhibited intermediate T2D enrichment and HbA1c values near the pre-diabetic range, while cluster 7 met T2D criteria. The remaining clusters, including the polyhormonal cluster 6, were classified as ND (Figures 2G-2I). Alpha cells showed no comparable pre-T2D intermediate state (Figure S3F). Spatial proximity between beta-cell clusters 2 and 7 in the integrated embedding suggested progressive transcriptional remodeling from early to advanced disease-associated states (Figure 2I).

Slingshot-based trajectory inference, rooted at cluster 3 based on its enrichment for young, lean ND donors, resolved three distinct beta-cell trajectories (Figure 2J). Lineage1 progressed toward an ND endpoint, Lineage2 transitioned through the pre-T2D cluster into a T2D-enriched endpoint, and Lineage3 diverged toward the polyhormonal ND cluster. A unified pseudotime representation enabled direct comparison across lineages.

INS expression remained largely stable along Lineage1, declined modestly along Lineage2, and showed the most pronounced reduction along Lineage3, despite the ND-enriched clinical profile of its endpoint (Figure 2K). Together, these results indicate that beta cells traverse multiple, structured transcriptional trajectories associated with distinct disease and metabolic contexts, whereas a comparable state diversification is limited or not readily detectable in alpha cells.

### Beta-cell aging is associated with ER-stress-biased and quiescent transcriptional trajectories independent of glycemic status

To resolve lineage-specific transcriptional programs underlying beta-cell aging, we re-analyzed Lineage1 at higher resolution. Cross-lineage comparison of highly variable genes (HVGs) revealed limited overlap between beta-cell lineages, with only 844 genes (24%) shared across all three (Figure S4 and table S5), indicating substantial lineage-specific transcriptional divergence.

Re-clustering Lineage1 beta cells (n = 16,733) using lineage-specific HVGs resolved four subclusters (Figures 3A-3C). One subcluster (C2) was enriched for beta cells from younger donors, whereas the remaining clusters were predominantly derived from older donors with higher BMI. All subclusters exhibited a below diabetic range median HbA1c values, indicating that this trajectory reflects aging-associated remodeling independent of overt hyperglycemia (Figures S5A).

**Figure 3:**
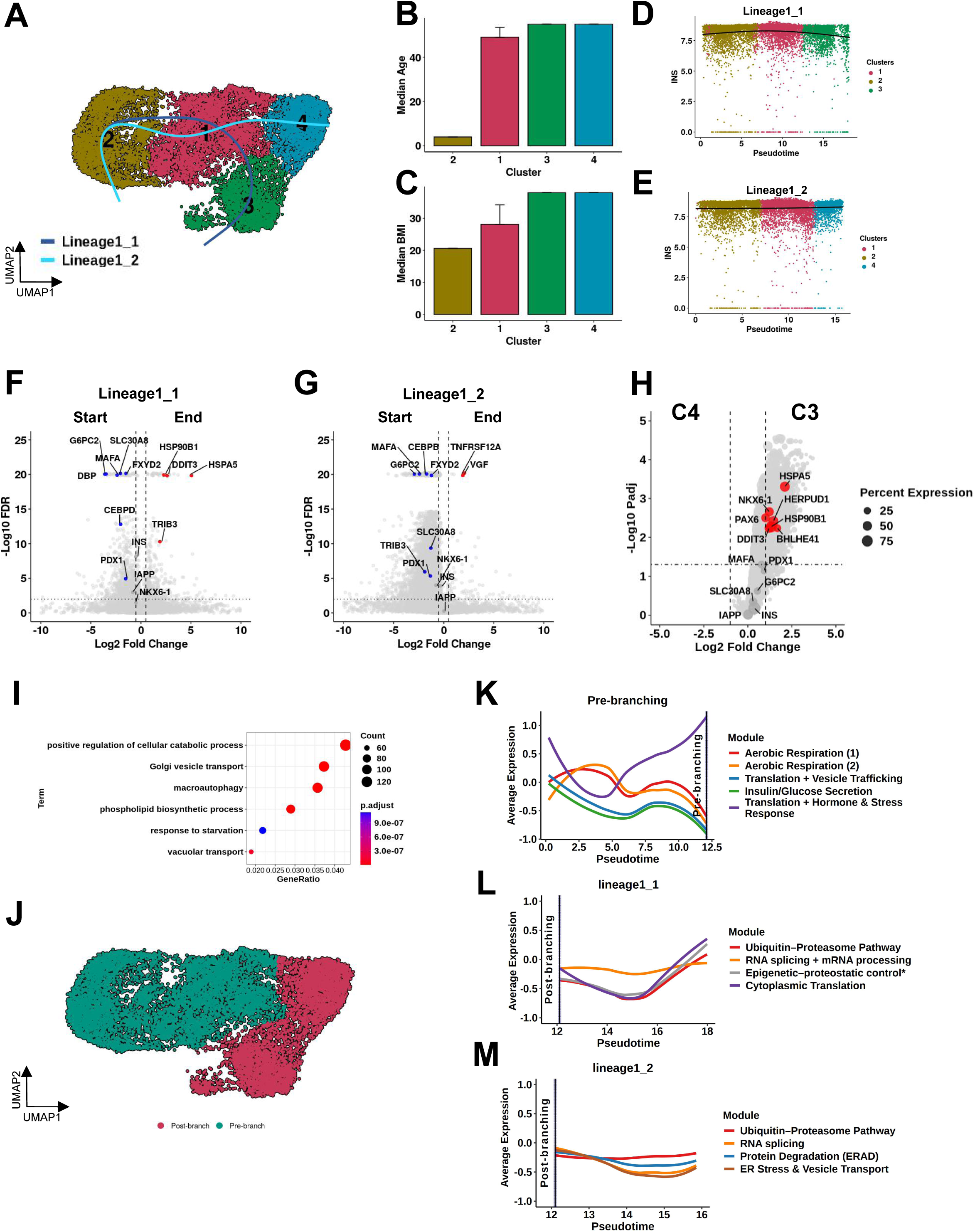
Lineage1 trajectory delineates the transition of human beta-cells in aging. (A) UMAP of beta-cells in part of the Lineage1 trajectory post re-clustering. The dark blue and light blue lines represent the inferred sub-lineages within Lineage1 beta-cells. (B) Median age and (C) BMI of the beta-cell clusters. Error bars represent the MAD. Scatter plot and smoother of *INS* expression changes along pseudotime for (D) Lineage1_1 and (E) Lineage1_2 sub-lineage trajectory. Volcano plots from diffEndTest comparing the terminal trajectories of (F) Lineage1_1 and (G) Lineage1_2. (H) Volcano plot of pseudobulk DEGs between clusters C3 and C4. Selected genes significantly upregulated (red) or downregulated (blue) are labeled. Dot sizes represent the percentage of cells expressing the gene. (I) Dotplot of selected GO BP terms enriched from C3 vs C4 pseudobulk DEGs analysis. (J) UMAP of beta-cells part of the Lineage1 trajectory showing pre- and post-branching. Pseudotime-resolved gene module dynamics along divergent beta-cell trajectories for (K) pre-branching, (L) in Lineage1_1, and (M) in Lineage1_2. “*” No GO terms reached FDR <0.2; annotation reflects the dominant theme of enriched terms.

Trajectory inference rooted at C2 identified two distinct aging-associated sub-trajectories (Lineage1_1 and Lineage1_2; Figures 3A). INS expression remained largely stable along both trajectories (Figures 3D and 3E), yet the endpoints diverged transcriptionally. Pseudotime analysis identified 2,006 genes changing along Lineage1_1 and 3,915 along Lineage1_2 (Figures 3F-3G and Table S6-S7).

Functional enrichment revealed that Lineage1_1 was characterized by induction of ER-stress, apoptotic, and proteostasis-related pathways, accompanied by repression of membrane potential and excitability programs (Figure S5B). In contrast, Lineage1_2 lacked enrichment for stress-response pathways but showed similar attenuation of excitability-associated genes, consistent with a transcriptionally quiescent state rather than an activated stress response (Figure S5C). Glucose-responsive genes, including *ABCC8, CACNA1C, PCSK1*, and *PDX1*, were preferentially reduced in the Lineage1_2 endpoint, whereas only *PCSK1* showed comparable reduction along Lineage1_1 (Figure S5D).

Pseudobulk differential expression analysis comparing lineage endpoints revealed preservation of core beta-cell identity markers (*INS, IAPP, G6PC2, MAFA*), with selective reduction of *NKX6-1* and *PAX6* (Figure 3H and Table S8). Genes enriched in the Lineage1_1 endpoint were associated with proteasomal processing, vesicle trafficking, autophagy, and unfolded protein response pathways (Figure 3I), accompanied by elevated expression of canonical stress markers including *HSPA5, HSP90B1*, and *DDIT3*.

Pseudotime-resolved module analysis revealed a shared early phase across both aging trajectories marked by downregulation of metabolic and insulin-secretion modules, followed by transient activation of translation-related programs (Figures 3J-3K). After bifurcation, Lineage1_1 exhibited reactivation of translation, RNA processing, and proteostatic control modules, alongside induction of ER-stress and senescence-associated genes. In contrast, Lineage1_2 showed minimal post-branching remodeling, consistent with stabilization in a quiescent, low-excitability state (Figures 3L-3M).

Together, these analyses indicate that beta-cell aging involves early functional attenuation followed by divergence into two transcriptional outcomes: one characterized by activation of ER-stress and proteostatic programs, and another marked by relative transcriptional quiescence, despite similar glycemic profiles.

### ER-stress-biased and exocrine-like transcriptional programs define divergent beta-cell trajectories associated with T2D

To characterize diabetes-associated beta-cell remodeling, we examined Lineage2 (13,853 beta cells), which was enriched for T2D donors. Re-clustering using lineage-specific features resolved five subpopulations showing a stepwise increase in T2D enrichment and HbA1c values, identifying a pre-diabetic-enriched cluster (C3) and two T2D-associated endpoints (C2 and C5) (Figures 4A-4C and S6A-S6D).

**Figure 4:**
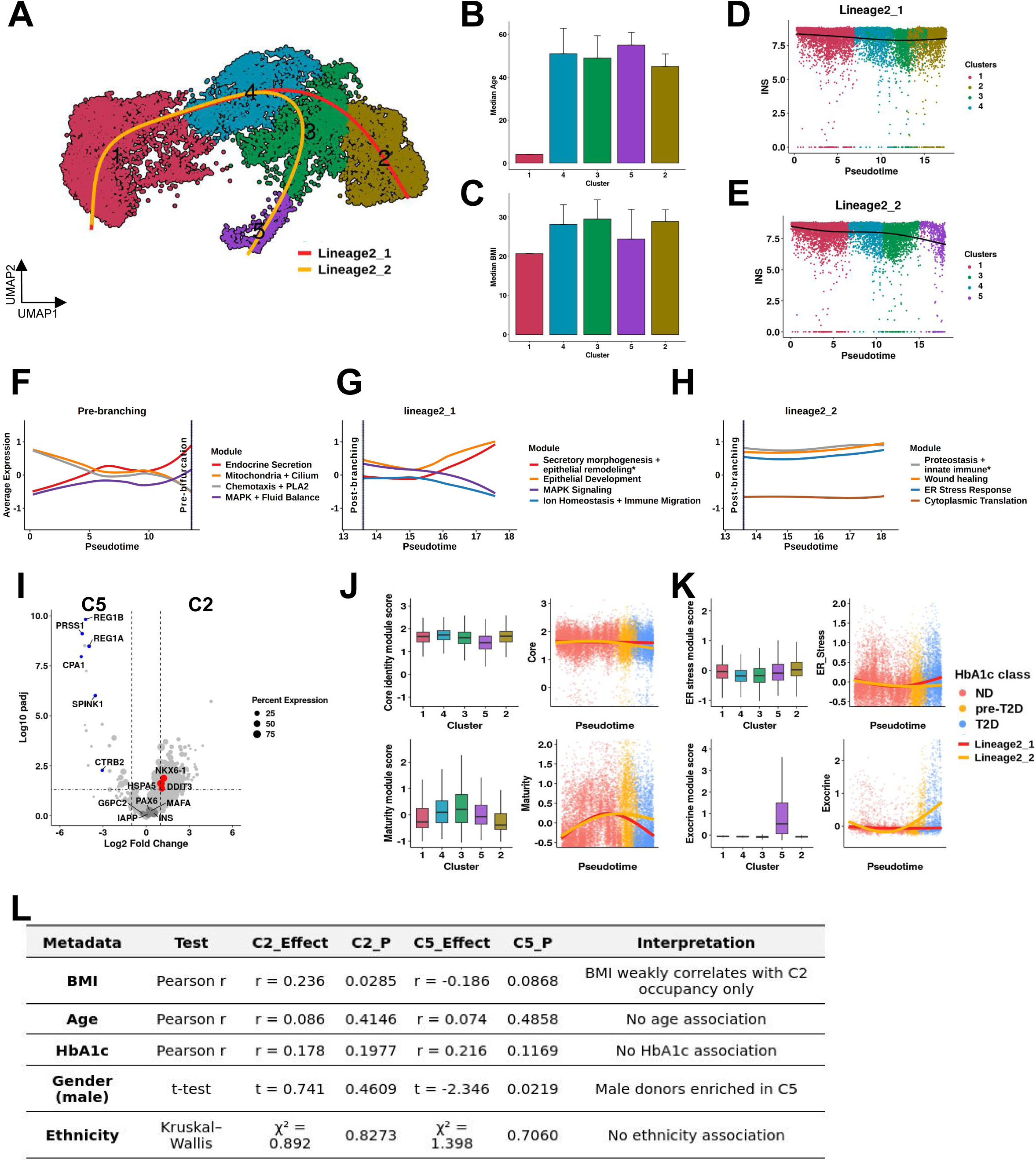
Lineage2 trajectory analysis reveals beta-cell states underlying T2D transitions. **(A)** UMAP of beta-cells part of the Lineage2 trajectory post re-clustering. The red and yellow lines represent the inferred sub-lineages within Lineage2 beta-cells. (B) Median age and (C) BMI of the beta-cell clusters. Error bars represent the Median Absolute Deviation (MAD). Scatter plot and smoother of INS expression changes along pseudotime for (D) Lineage2_1 and (E) Lineage2_2 sub-lineage trajectories. Pseudotime-resolved gene module dynamics in (F) pre-branching, (G) Lineage2_1, and (H) Lineage2_2. (I) Volcano plots of pseudobulk DEGs comparing clusters C2 vs C5. Selected genes significantly upregulated (red) or downregulated (blue) are labeled. Dot sizes represent the percentage of cells expressing the gene. Module score comparisons for (J) Core/Maturity and (K) ER Stress/Exocrine program module scores between Lineage2_1 and Lineage2_2. Dots are colored by HbA1c classification (ND, pre-T2D, T2D), and smoothed lines represent LOESS regression fits. Lineage assignments are shown in the legend. Module scores were computed using Seurat’s AddModuleScore function. (L) Donor-level associations with Lineage2 endpoint clusters. Effect sizes and P values are shown for clusters C2 and C5 using Pearson correlation (BMI, Age, HbA1c), t-test (gender), and Kruskal–Wallis test (ethnicity). BMI was weakly associated with C2 occupancy, and male donors were enriched in C5; other variables showed no significant associations. “*” No GO terms reached FDR < 0.2; annotation reflects the dominant theme of enriched terms.

Trajectory inference revealed that Lineage2 bifurcates from C3 into two distinct diabetes-associated paths: Lineage2_1 terminating in C2 and Lineage2_2 terminating in C5 (Figures 4A). INS expression declined modestly along Lineage2_2 but remained comparatively stable along Lineage2_1, indicating trajectory-specific remodeling of beta-cell functional programs (Figures 4D and 4E).

Pseudotime-resolved module analysis revealed shared early transcriptional changes followed by divergence into two distinct adaptive states (Figures 4F-4H). Prior to bifurcation, beta cells upregulated secretory and MAPK-related modules while repressing mitochondrial and ciliary programs. After branching, Lineage2_1 showed sustained activation of exocytosis and epithelial-development modules, consistent with a proteostatic, secretory-stress-associated state. In contrast, Lineage2_2 progressively enriched ER-stress/proteostasis, wound-healing, and translation-associated modules, reflecting a distinct adaptive response.

Pseudobulk endpoint analysis further distinguished the two trajectories. The C2 endpoint exhibited higher expression of canonical ER-stress genes, including *HSPA5* and *DDIT3*, whereas C5 was marked by strong induction of exocrine-associated transcripts (*PRSS1, REG1A/B, CPA1, SPINK1*) and downregulation of beta-cell identity factors such as *NKX6-1* and *PAX6* (Figure 4I and Table S9). Although only a subset of ER-stress genes reached significance at the endpoint, their overall enrichment was biased toward C2, consistent with sustained proteostatic stress along Lineage2_1.

Pathway analysis of pseudotime-associated genes reinforced these distinctions. Lineage2_1 was enriched for oxidative stress, immune signaling, and differentiation pathways, whereas Lineage2_2 showed enrichment for secretion-related, lipid catabolic, hormone-response, and transcriptional activation programs (Figures S6E-S6G and Tables S10-S11).

Consistent with these transcriptional profiles, beta-cell identity module scores indicated that C2 retained core identity features with reduced maturity, while C5 preserved maturity-associated programs but showed stronger enrichment of exocrine-like transcriptional signatures (Figures 4J-4K).

Donor-level analysis revealed modest associations with clinical traits. Body mass index correlated with occupancy of the proteostatic C2 state, whereas the exocrine-like C5 state was enriched in male donors and associated with elevated integrated stress response signaling, including eIF2α pathway components, consistent with translational stress downstream of ER perturbation (Figures 4L and S6H). Age, HbA1c, and ethnicity were not significantly associated with either endpoint.

Together, these results indicate that T2D-associated beta-cell remodeling proceeds along two distinct transcriptional trajectories emerging from a shared pre-diabetic state: one characterized by sustained proteostatic stress with partial loss of maturity, and another marked by exocrine-like transcriptional reprogramming accompanied by erosion of core beta-cell identity.

### Lipid-associated metabolic stress defines an early beta-cell state with poly-hormonal transcriptional features

Lineage3 (n = 8,882 beta cells) resolved into three subclusters (C1-C3) differing primarily by donor metabolic status rather than glycemia (Figures 5A-5C). C1 was enriched for beta cells from young, lean donors, whereas C2 and C3 were dominated by cells from older, overweight or obese donors. C3 showed the highest enrichment of T2D-associated cells within Lineage3 (36.3%), although median HbA1c values remained below the diagnostic threshold for diabetes (Figures S7A-S7C).

**Figure 5:**
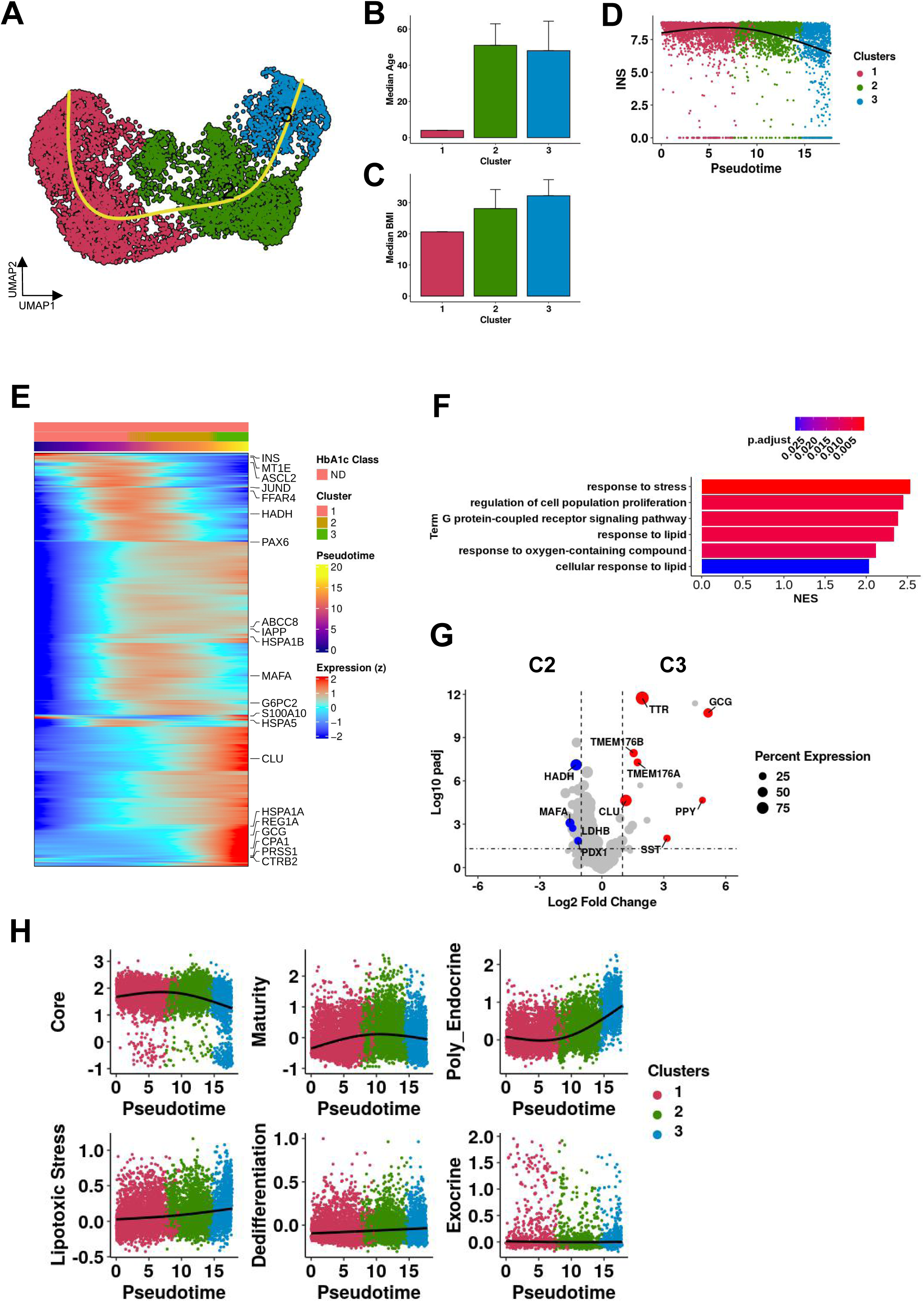
Lineage3 trajectory delineates the transition of beta-cells from cluster 1 to cluster 3. (A) UMAP of beta-cells part of the Lineage3 trajectory post re-clustering. (B) Median age and (C) BMI of the beta-cell clusters. Error bars represent the MAD. (D) Scatter plot and smoother of *INS* expression changes along pseudotime. (E) Trajectory heatmap showing gene expression along pseudotime. (F) GSEA of genes positively associated with Lineage3 pseudotime trajectory using GO BP terms. (G) Volcano plot of pseudobulk DEGs comparing C3 vs C2. Selected genes significantly upregulated (red) or downregulated (blue) are labeled. Dot sizes represent the percentage of cells expressing the gene. (H) Module score trends across pseudotime for key beta-cell identity, stress, and transdifferentiation programs. LOESS fits highlight decline of Core/Maturity markers and induction of poly-hormonal and lipotoxic stress signatures in C3.

Slingshot trajectory inference rooted in C1 revealed progressive functional remodeling toward the C3 endpoint, marked by a sharp decline in INS expression along pseudotime (Figures 5D-5E and Table S12). Gene set enrichment analysis of pseudotime-associated genes identified lipid metabolism, cellular stress responses, and GPCR signaling as dominant programs along this trajectory, with comparatively weaker enrichment of glucotoxic stress pathways (Figure 5F). Consistently, pseudobulk comparison of C3 versus C2 confirmed downregulation of core beta-cell functional genes (*IAPP, MAFA, PDX1*) alongside induction of stress-associated chaperones (*CLU, TMEM176A, TMEM176B*) (Figure 5G and Table S13).

A defining feature of the C3 state was increased expression of multiple endocrine hormone transcripts, including *GCG, PPY*, and *SST*, consistent with a poly-hormonal transcriptional profile evident at both pseudobulk and single-cell resolution (Figures 2F and S7D). Module score analysis showed marked decay of core beta-cell transcriptional programs at late pseudotime, a transient rise in maturity signatures in C2, and strong induction of poly-hormonal endocrine modules toward C3, while exocrine and progenitor-like programs remained unchanged (Figure 5H and S7D). These patterns indicate endocrine identity remodeling under lipid-associated stress rather than exocrine transdifferentiation or progenitor reversion.

At the donor level, C3 occupancy correlated strongly with BMI (r = 0.375, p = 0.002), but not with age, sex, or HbA1c (Figure S7E). Ethnicity was also significantly associated with variation in C3 representation (p = 0.019). Together, these findings define Lineage3 C3 as an early beta-cell state driven by lipid-associated metabolic stress and characterized by poly-hormonal transcriptional remodeling that occurs independently of overt hyperglycemia.

### Divergent diabetes-associated exocrine-like remodeling and lipid-associated dedifferentiation underlie beta-cell identity loss

Our integrated atlas identified two beta-cell populations that undergo identity loss through distinct transcriptional routes under metabolic stress. Lineage2 cluster 5 (L2_C5), enriched in T2D donors, showed a moderate reduction in *INS* expression accompanied by induction of acinar-associated gene programs, including *CPA1* and *PRSS1/2*, consistent with exocrine-like transcriptional remodeling (Figures 4). In contrast, Lineage3 cluster 3 (L3_C3) exhibited a poly-hormonal endocrine state characterized by *GCG* and *SST* co-expression, pronounced *INS* downregulation, and enrichment for lipid-handling and stress-associated pathways (Figures 5). While L2_C5 cells retained key beta-cell identity factors such as *MAFA* and *PDX1* alongside exocrine gene induction, L3_C3 cells showed increased expression of alpha- and delta-lineage markers (e.g., *ARX, PAX6, SST, GCG*), indicating distinct molecular routes to beta-cell identity loss (Figures S8A).

To characterize the regulatory organization underlying these divergent transcriptional programs, we applied miEdgeR to construct gene–gene association networks based on mutual information (MI), which captures both linear and non-linear relationships in single-cell expression data. Gene expression values were discretized using adaptive binning to improve robustness in sparse single-cell profiles, and network construction was restricted to the top 5% of MI edges to focus on the strongest associations. Network structure was stabilized using repeated edge subsampling (percolation), and gene communities were identified using label-propagation clustering. Because genes can contribute to more than one biological process, community assignments were expanded based on neighborhood overlap, allowing regulatory modules to share genes and capture graded regulatory structure (Figure 6A).

**Figure 6:**
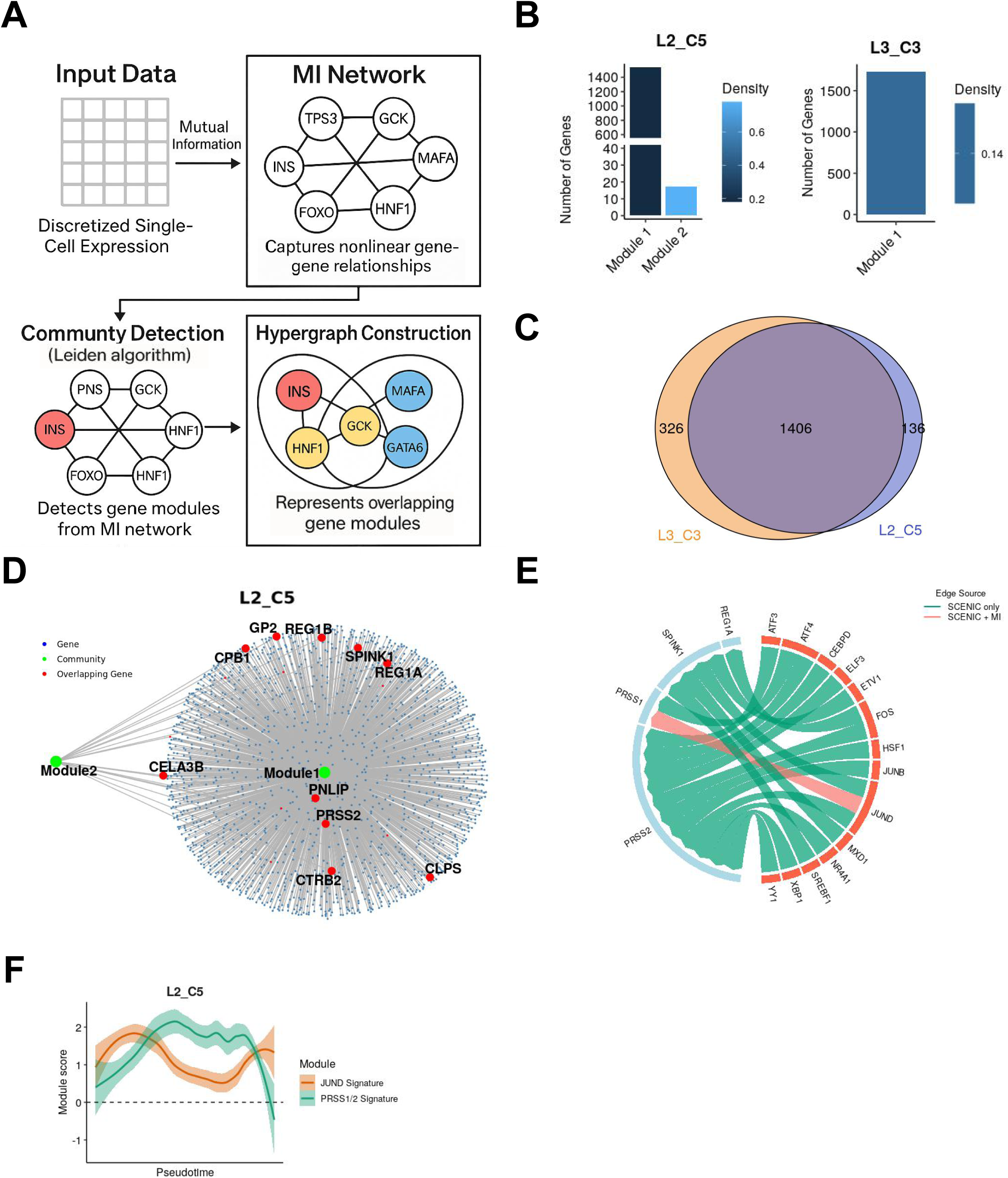
Divergent mutual information networks reveal distinct regulatory circuits driving beta-cell identity loss in L2_C5 and L3_C3. (A) Schematic of the miEdgeR framework. Single-cell expression is discretized and pairwise mutual information (MI) is computed to infer gene-gene association networks. High-confidence edges are retained using consensus filtering, and overlapping gene modules are identified by label-propagation clustering with neighborhood overlap. (B) Module size and density for MI-derived networks in L2_C5 and L3_C3. (C) Overlap between stress-associated modules in L2_C5 and L3_C3. (D) MI network structure in L2_C5 showing stress-associated and exocrine-like modules with shared genes. (E) SCENIC-predicted transcription factor–target interactions in L2_C5, with links supported by MI network connectivity highlighted. (F) Module-score trajectories along L2_C5 pseudotime showing earlier activation of JUND-associated signatures relative to PRSS1/2 expression.

In L2_C5, this approach resolved two partially overlapping programs: a large stress-associated module containing heat-shock and unfolded-protein-response genes, and a compact exocrine-like module characterized by digestive enzyme transcripts forming a tightly co-regulated subnetwork (Figures 6B and S8B). In L3_C3, miEdgeR identified a dominant stress-associated module containing endocrine and stress-response genes, consistent with a dedifferentiated poly-hormonal state rather than exocrine-like remodeling (Figures 6B and S8B). Comparison of stress-associated modules between L2_C5 and L3_C3 revealed a substantial shared gene set defining a conserved stress core, alongside cluster-specific genes reflecting either exocrine-like remodeling in L2_C5 or lipid-associated endocrine dedifferentiation in L3_C3 (Figure 6C and Table S14).

In L2_C5, overlap between stress-associated and exocrine-like modules identified a subset of genes shared between both programs. Within the exocrine-like module, highly connected genes were distinguished from more weakly connected overlapping genes, which were classified as transitional (Figure 6D).

Integration with SCENIC enabled annotation of transcription factor–target interactions supported by both MI network connectivity and motif-based inference. Among transcription factors linked to exocrine-associated genes, JUND was consistently supported by both approaches, appearing as a central hub in the MI-derived stress-associated module and as a predicted regulator of multiple exocrine-associated targets in SCENIC (Figure 6E). Consistent with this network positioning, JUND module scores increased earlier along the L2_C5 pseudotime axis than PRSS1/2 exocrine gene scores (Figure 6F), indicating earlier engagement of stress-associated signatures relative to exocrine-like output genes.

## DISCUSSION

Here, we assembled a unified atlas of 266,614 human pancreatic cells by integrating 18 publicly available scRNA-seq datasets spanning diverse technologies, donor demographics, and disease states. By minimizing study- and platform-specific effects, this integration recovered all major pancreatic cell types and resolved beta-cell transcriptional states associated with aging and metabolic stress. The atlas provides a consolidated reference for interpreting human islet heterogeneity using public single-cell data.

Trajectory-based analyses identified three major beta-cell trajectories linked to aging, diabetes-associated stress with exocrine-like remodeling, and lipid-associated metabolic stress. These trajectories do not represent fixed lineage relationships, but instead describe structured patterns of beta-cell state remodeling observed across individuals. In contrast to the relative stability of alpha-cell states, beta cells showed substantial transcriptional plasticity along all three trajectories, often preceding marked reductions in INS expression.

The aging-associated trajectory (Lineage1) diverged under normoglycemic conditions into two outcomes: one marked by activation of ER stress and senescence-related programs, and another characterized by transcriptional quiescence with reduced excitability and insulin-secretion machinery in the absence of overt stress signaling. These findings indicate that aging-related beta-cell functional decline can proceed through multiple transcriptional routes without immediate loss of core identity.

The diabetes-associated trajectory (Lineage2) diverged from a shared pre-diabetic intermediate into two distinct outcomes. One path retained core beta-cell identity while engaging sustained proteostatic and ER stress programs, whereas the other showed erosion of beta-cell identity accompanied by induction of exocrine-associated transcriptional programs. Both outcomes emerged from the same precursor state, indicating that beta-cell dysfunction in T2D is not a single linear process but reflects alternative stress-adaptive transcriptional responses.

A third trajectory (Lineage3) captured a lipid-associated metabolic stress state characterized by poly-hormonal endocrine features and pronounced INS downregulation, occurring largely under normoglycemic conditions. This state was strongly associated with adiposity and was not accompanied by exocrine transdifferentiation or progenitor-like reversion, suggesting that lipid-associated stress is linked to remodeling of endocrine identity without evidence for classical dedifferentiation.

Donor-level analyses indicated that beta-cell transcriptional outcomes are influenced by systemic context. BMI correlated with lipid-associated beta-cell states, whereas sex differences were evident in T2D-associated metabolic stress trajectories. Associations with ethnicity likely reflect complex genetic, metabolic, and environmental factors and should be interpreted cautiously.

This study is limited by its reliance on transcriptomic inference and cross-sectional sampling, which precludes causal conclusions. Residual technical variability and incomplete transcript capture inherent to scRNA-seq data cannot be fully excluded. Independent validation across additional cohorts will be required to assess generalizability.

In summary, this integrated atlas shows that human beta-cell dysfunction is not uniform but instead involves multiple, transcriptionally distinct stress-associated routes. By resolving aging-, T2D-, and lipid-associated beta-cell trajectories, this work provides a systems-level framework for understanding beta-cell plasticity and heterogeneity in diabetes.

## LIMITATIONS

While this study provides an integrative view of human beta-cell transcriptional heterogeneity, several limitations should be noted. Lineage assignment and pseudotime reconstruction rely on computational inference applied to cross-sectional single-cell transcriptomic data. Accordingly, the trajectories described here represent descriptive patterns of transcriptional state variation rather than direct lineage relationships or causal models of disease progression. In addition, the resolution of inferred trajectories is constrained by the availability and consistency of donor metadata, including HbA1c and body mass index. All analyses are based on transcriptomic measurements and therefore do not directly assess protein abundance, post-translational regulation, or cellular function.

Integration of datasets generated across multiple platforms and protocols introduces unavoidable technical heterogeneity despite extensive quality control and batch correction. Residual technical effects, incomplete transcript capture, and dropouts inherent to scRNA-seq data may influence detection of low-abundance genes and subtle regulatory signals. In addition, the focus on protein-coding genes precludes assessment of non-coding and epigenetic regulatory mechanisms.

Finally, classification of beta-cell states using clinical thresholds such as HbA1c simplifies the heterogeneous nature of diabetes. Genetic background, treatment history, environmental exposures, and temporal disease dynamics are incompletely represented in available datasets. Future studies incorporating longitudinal sampling, multi-omic profiling, and richer clinical annotation will be required to further refine the transcriptional frameworks described here.

## ACKNOWLEDGEMENTS

We would like to thank the investigators and consortia who generated and made publicly available the human pancreatic single-cell RNA-sequencing datasets used in this study. Computational analyses were supported by the NUS Information Technology Research Computing group under grant NUSREC-HPC-00001.

## AUTHOR CONTRIBUTIONS

E.I. conceived the study, performed all analyses, developed the computational methods, generated visualizations, and wrote the manuscript.

## METHODS

### Data acquisition and preprocessing

Raw sequencing data from 18 publicly available human pancreatic scRNA-seq datasets were obtained from GEO and ENA (Table S1) and uniformly reprocessed from raw reads to minimize cross-study technical variability. Droplet-based datasets were processed using Cell Ranger (v6.1.2), while full-length and plate-based datasets were aligned to GRCh38 using STAR, followed by gene-level quantification with featureCounts or UMI-tools as appropriate. All datasets were converted to sparse gene-cell count matrices and processed using a unified downstream workflow.

Ambient RNA contamination in droplet-based datasets was corrected using SoupX. Low-quality cells were filtered based on dataset-specific thresholds for detected genes, mitochondrial content, and transcriptomic complexity. Analyses were restricted to protein-coding genes. Doublets were inferred using DoubletFinder and scDblFinder, and cells identified by either method were excluded prior to integration.

All analyses were performed on publicly available scRNA-seq datasets obtained from GEO and ENA. No unpublished experimental data or proprietary datasets were used. Processed intermediate objects were regenerated from public sources for the purposes of this study.

### Global integration and clustering

Following quality control, datasets were merged and normalized in Seurat (v4.3.0.1). Highly variable genes were identified, data were scaled while regressing out library size and mitochondrial content, and batch correction was performed using Harmony with study identity as the integration variable. Clustering was performed using Leiden community detection on Harmony-corrected principal components, and UMAP was used for visualization.

### Cell-type annotation and validation

Cell-type identities were assigned based on canonical marker expression and independently validated using a supervised scPred classifier trained on well-annotated reference datasets. Classifier predictions were summarized at the cluster level and used to support marker-based annotations.

### Lineage-specific re-integration and trajectory inference

For lineage analyses, beta and alpha cells were subsetted and re-integrated using SCTransform to enhance resolution of continuous transcriptional variation. Monocle3 was used for low-dimensional embedding and clustering. Trajectories and pseudotime were inferred using Slingshot without imposing a predefined hierarchy. Pseudotime-associated genes were identified using tradeSeq, with generalized additive models fitted to expression dynamics along each lineage.

### Pseudobulk differential expression

Raw counts were aggregated into pseudobulk samples by donor, cluster, and lineage. Genes expressed in at least 5% of cells were retained. Differential expression analysis was performed using DESeq2, with size factors proportional to contributing cell counts to correct for unequal aggregation.

### Module detection and trajectory analysis

Co-regulated gene modules were identified using a custom PCA-based approach applied to binned pseudotime expression matrices. Modules were defined using hierarchical clustering with dynamic tree cutting and annotated via Gene Ontology enrichment. Predefined biological programs were quantified using Seurat’s AddModuleScore and visualized across pseudotime.

### Statistical analysis and reproducibility

Multiple testing correction was performed using the Benjamini-Hochberg method. All analyses used fixed random seeds.

### Mutual information network inference and regulatory annotation

To characterize gene-gene association structure within selected beta-cell clusters, mutual information (MI) networks were inferred using the custom R package *miEdgeR*. For each cluster, genes expressed in at least 5% of cells were retained, and MI was computed for all gene pairs among the selected genes following adaptive discretization of expression values. Networks were constructed by retaining edges corresponding to the upper quantile of MI values, and robustness was improved using a percolation-based consensus procedure with repeated edge subsampling. Gene communities were identified using label-propagation clustering, allowing overlapping module structure. https://github.com/iichelhadi/miEdgeR.

Putative transcription factor–target relationships were inferred using SCENIC, and regulon activity was quantified using AUCell. SCENIC predictions were integrated with MI network structure to annotate transcription factor-target associations supported by both approaches. These analyses were used to contextualize regulatory architecture and are interpreted as descriptive rather than causal.

## Data Availability

All raw scRNA-seq datasets analyzed in this study are publicly available from their original repositories as cited.

The integrated and harmonized pancreas atlas (~266,000 cells), including metadata, annotations, and processed Seurat object used for analysis, has been deposited in Zenodo and is available at https://doi.org/10.5281/zenodo.18546206

## SUPPLEMENTARY FIGURE LEGENDS

**Figure S1:**
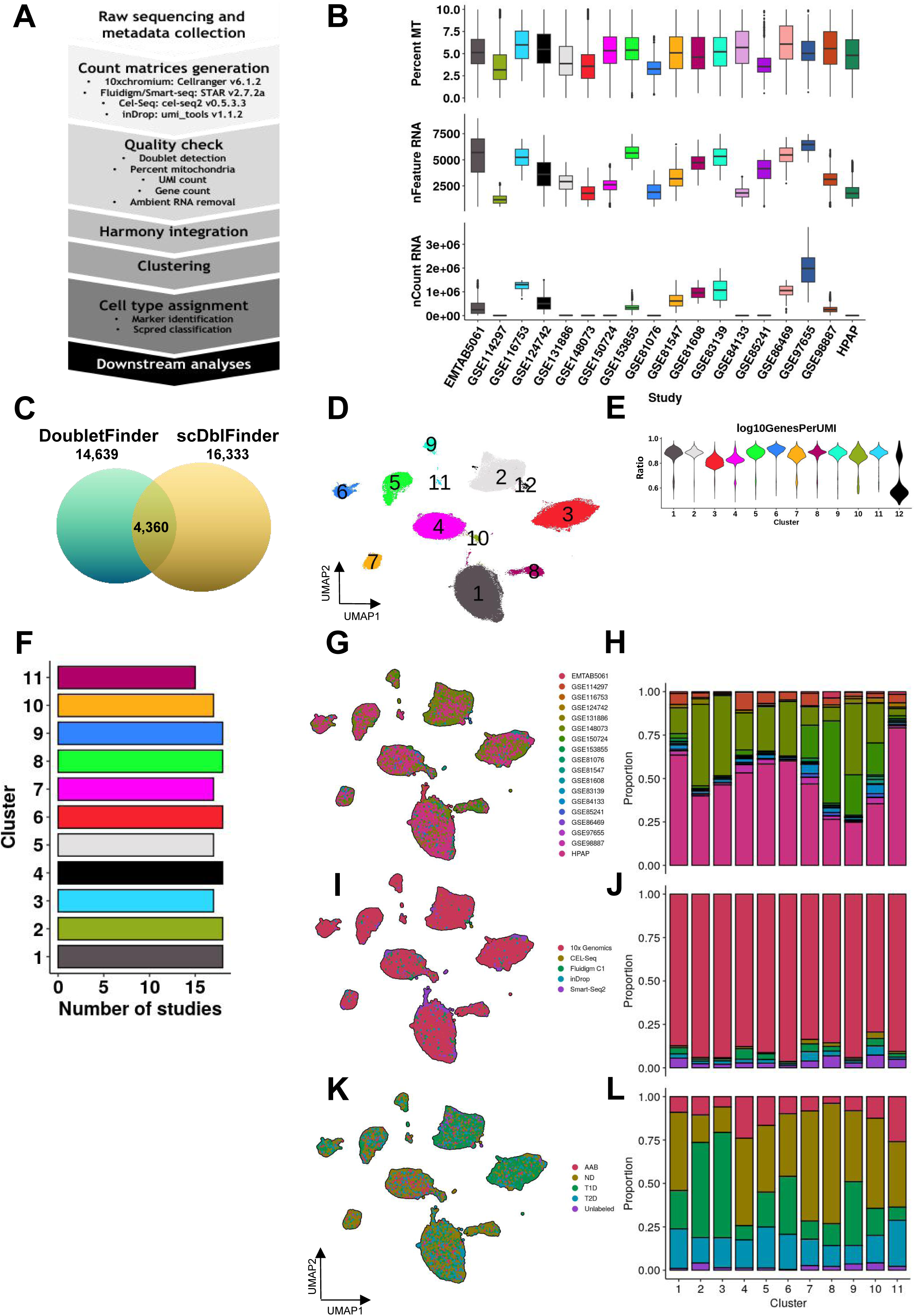
Characterization of 18 human pancreatic scRNA-seq studies across sequencing platform and disease conditions. (A) Overview of preprocessing, quality control, and integration steps used to construct the pancreatic scRNA-seq atlas. (B) Study-specific distributions of quality control metrics, including mitochondrial gene percentage (percent.mt), number of detected genes per cell (nFeature_RNA), and total UMI counts (nCount_RNA). (C) Venn diagram showing overlap of predicted doublets identified by DoubletFinder and scDblFinder. (D) UMAP of integrated cells following Harmony correction and clustering across all 18 studies. (E) Cluster-wise comparison of library complexity across established clusters. (F) Number of studies contributing cells to each cluster; cluster 12 was excluded due to outlier study-specific bias. (G–H) Study-wise distribution and contribution proportions per cluster. (I) Distribution and (J) proportion of sequencing platforms contributing to each cluster. (K) Distribution and (L) proportion of disease conditions (ND, AAB, T1D, T2D, unlabeled) contributing to each cluster.

**Figure S2:**
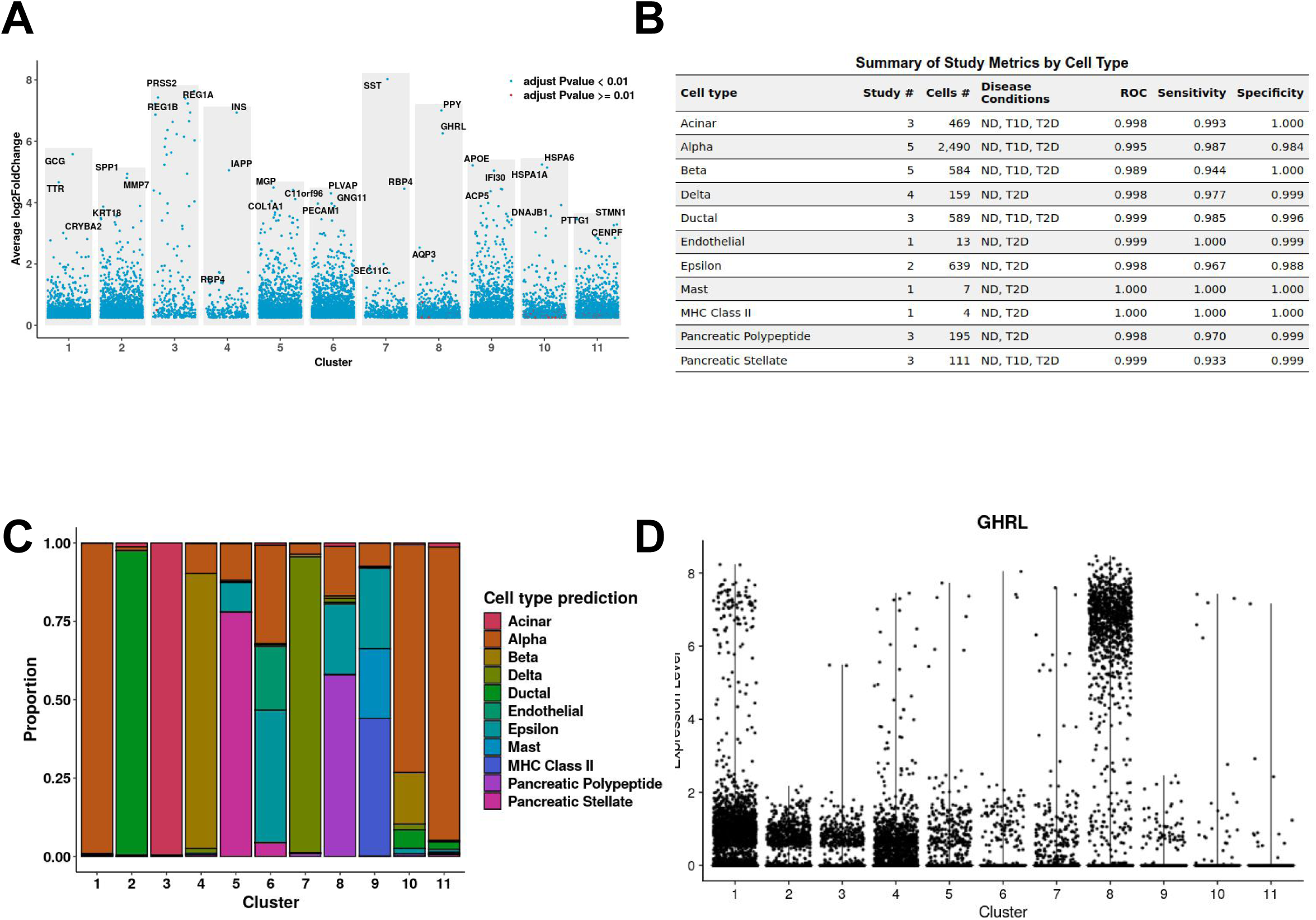
Pancreatic cell type identification and annotation using known markers and machine learning-trained classifier. (A) Manhattan plot showing scaled expression of genes enriched in each cluster. The top three upregulated and downregulated genes per cluster are labeled. (B) Summary table of the scPred classifier training, including number of studies contributing each annotated cell type, number of cells used for training, and classifier performance metrics (ROC AUC, sensitivity, specificity), along with disease condition distribution of training data. (C) Proportion of predicted cell types per cluster as inferred by the trained scPred classifier. (D) Violin plot of *GHRL* expression in identified clusters showing a subset of cluster 8 cells expressing high levels of the Epsilon cell marker.

**Figure S3:**
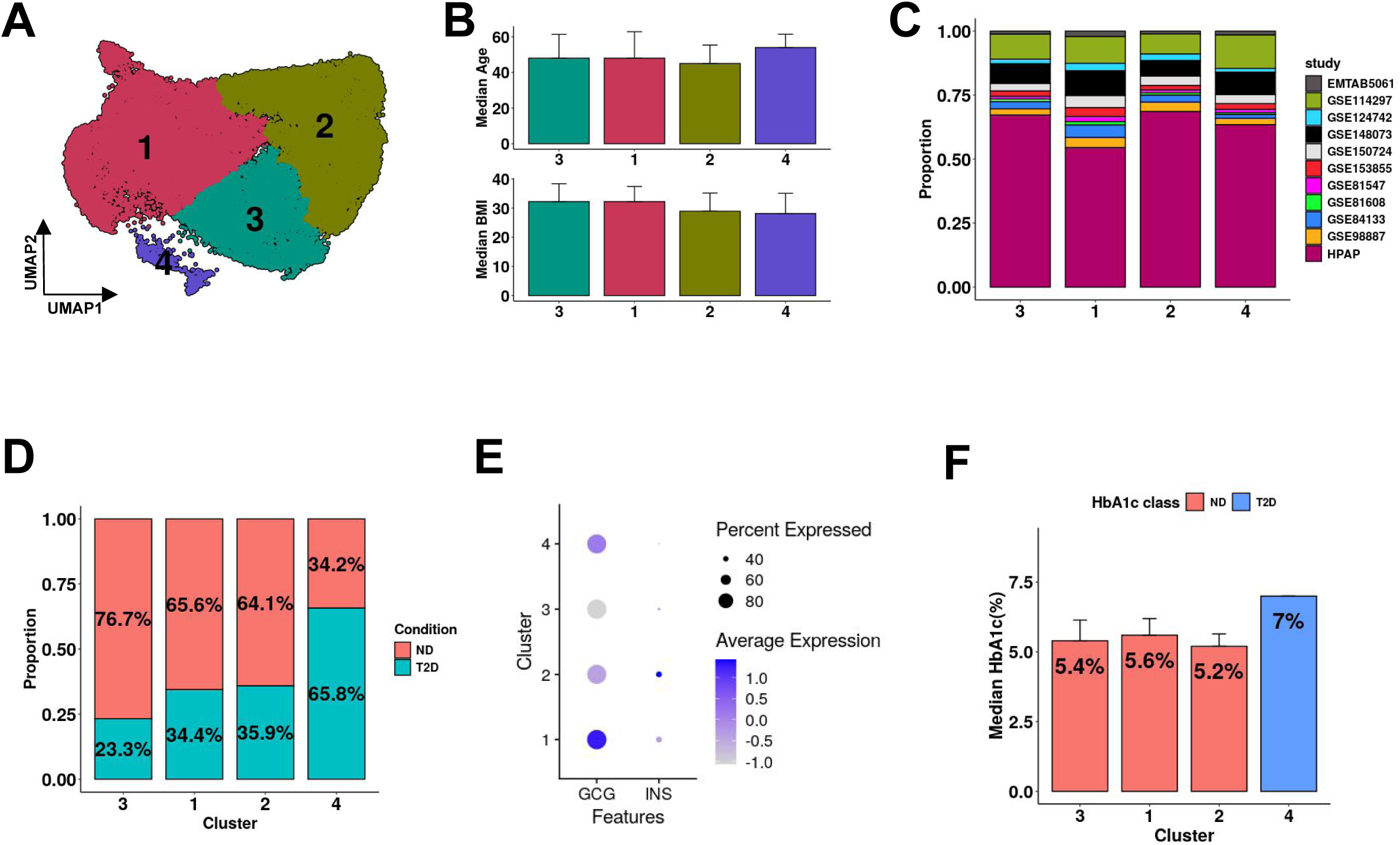
Integrated ND and T2D alpha-cells do not show transition to a dysfunctional state. (A) UMAP of SCTransform-integrated alpha-cells with clusters identified using Monocle3. (B) Median age and BMI of donors contributing to each alpha-cell cluster. Error bars represent the Median Absolute Deviation (MAD). (C) Proportion of individual studies and (D) ND and T2D donor cells in each alpha-cell cluster. (E) Dot plot of expression levels of *GCG* and *INS* across alpha-cell clusters. (F) Subclassification of alpha-cell clusters based on median HbA1c levels. Error bars represent the MAD.

**Figure S4:**
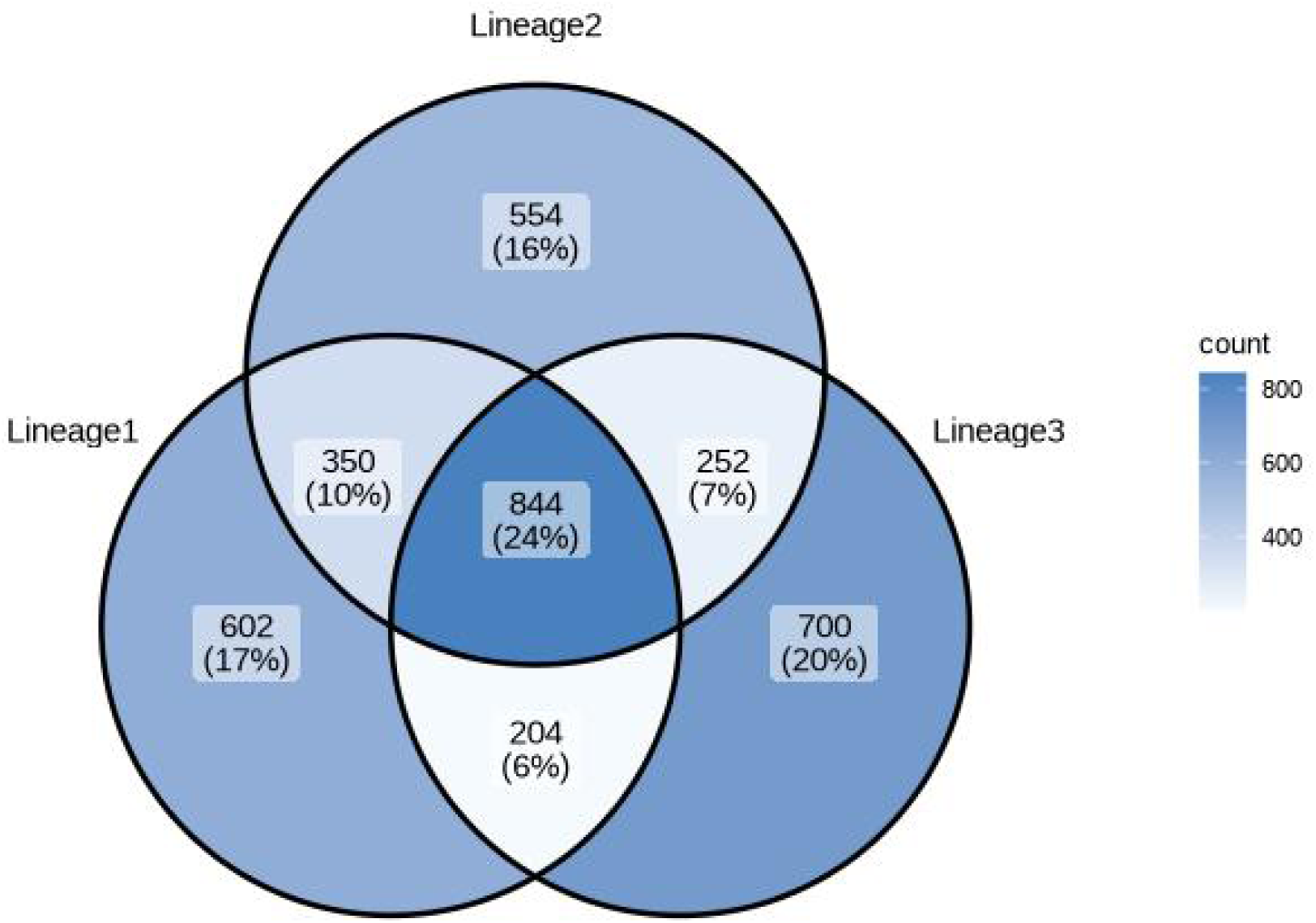
Overlap of highly variable genes (HVGs) across beta-cell lineages reveals lineage-specific transcriptional programs. Venn diagram showing the number of highly variable genes identified within each beta-cell lineage: Lineage1 (aging-associated), Lineage2 (T2D-enriched), and Lineage3. HVGs were calculated independently for each lineage using the FindVariableFeatures function after lineage-specific subsetting. Counts in the Venn diagram denote the number of genes unique or shared between one, two, or all three lineages.

**Figure S5:**
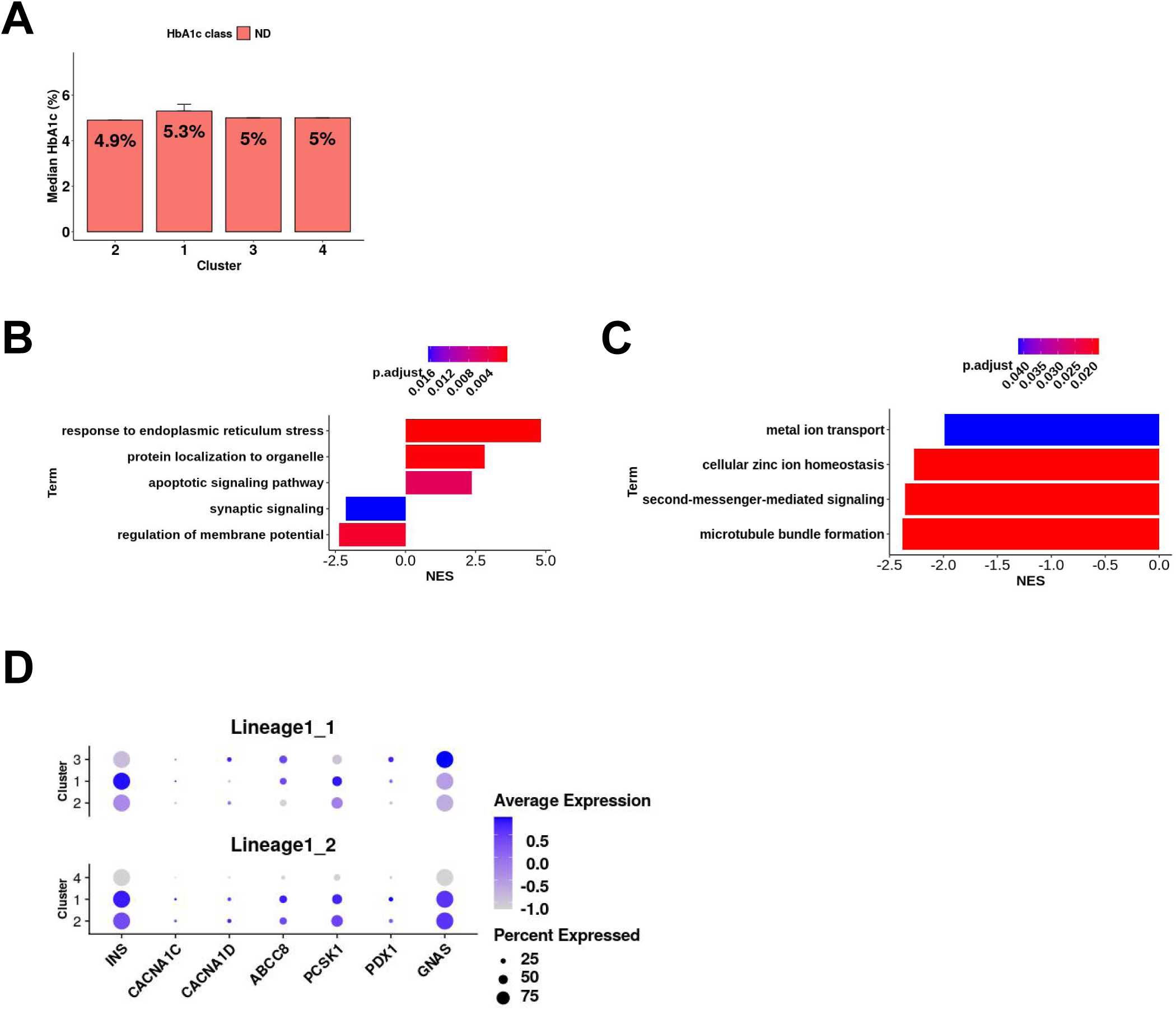
Lineage1 beta-cells reveal subcluster heterogeneity associated with ER stress and vesicle trafficking. (A) Barplot showing the median HbA1c across clusters. Error bars represent MAD. GSEA barplots of pseudotime-associated genes in (B) Lineage1_1 and (C) Lineage1_2. NES: normalized enrichment score. (D) Dotplot of glucose-responsive genes in Lineage1_1 vs Lineage1_2. Dot size indicates percent of cells expressing the gene; color indicates average scaled expression.

**Figure S6:**
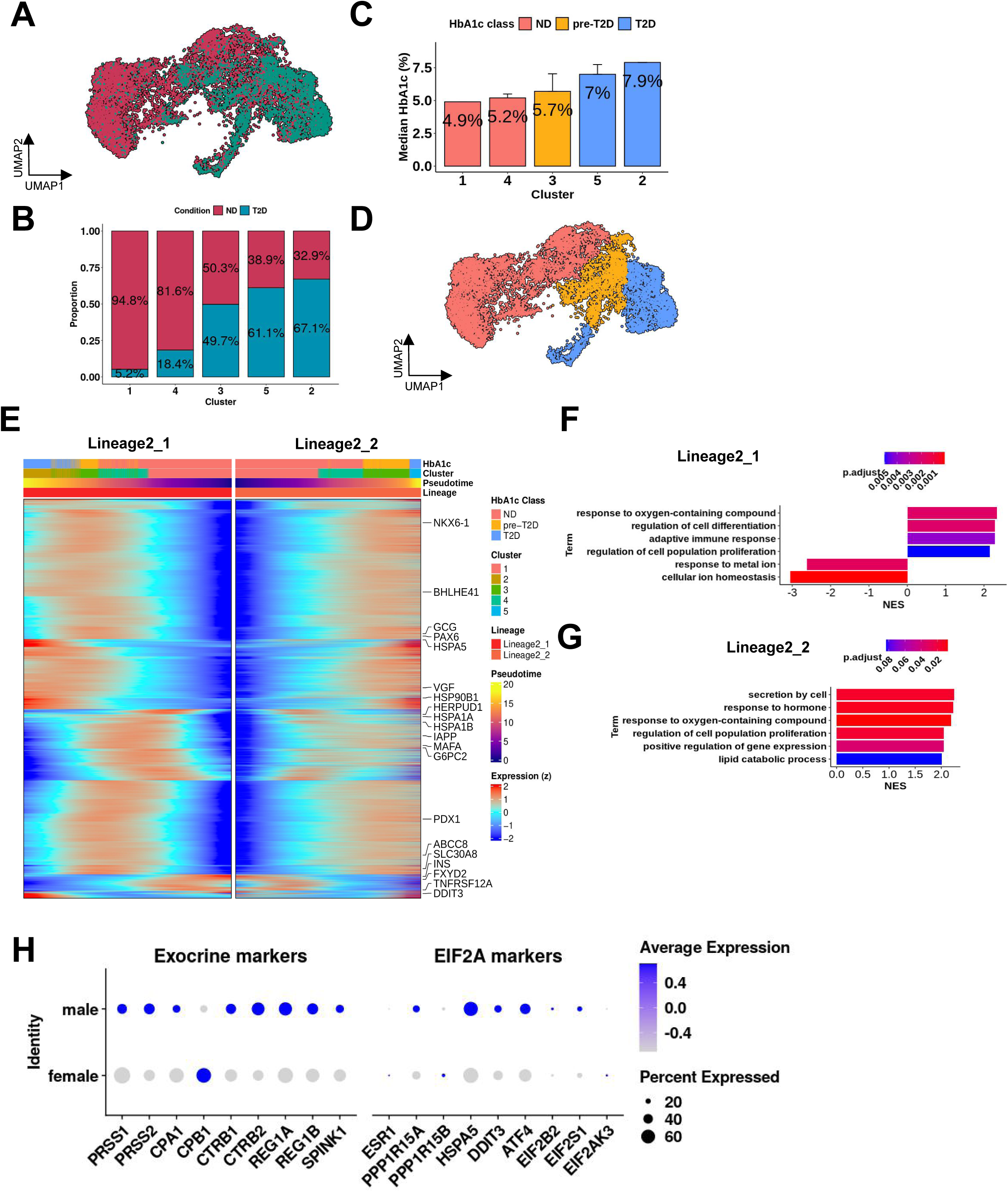
Divergent subcluster dynamics and sex bias along the Lineage2 trajectory. (A) UMAP colored by disease condition (ND vs T2D). (B) Proportion of ND and T2D cells per subcluster, highlighting progressive T2D enrichment from cluster 1 to cluster 2. (C) Barplot showing median HbA1c percentage per cluster. Error bars represent the Median Absolute Deviation (MAD). (D) UMAP of Lineage2 clusters colored by their median HbA1c classification. (E) Pseudotime-ordered heatmap of gene expression across Lineage2_1 (left) and Lineage2_2 (right). GSEA plots of pseudotime-associated genes in (F) Lineage2_1 and (G) Lineage2_2, respectively. (H) Dotplot of selected Exocrine and eIF2α pathway markers showing differential expression by sex (male vs female) within Lineage2 cluster 5. Dot size indicates percent of expressing cells; color represents average scaled expression.

**Figure S7:**
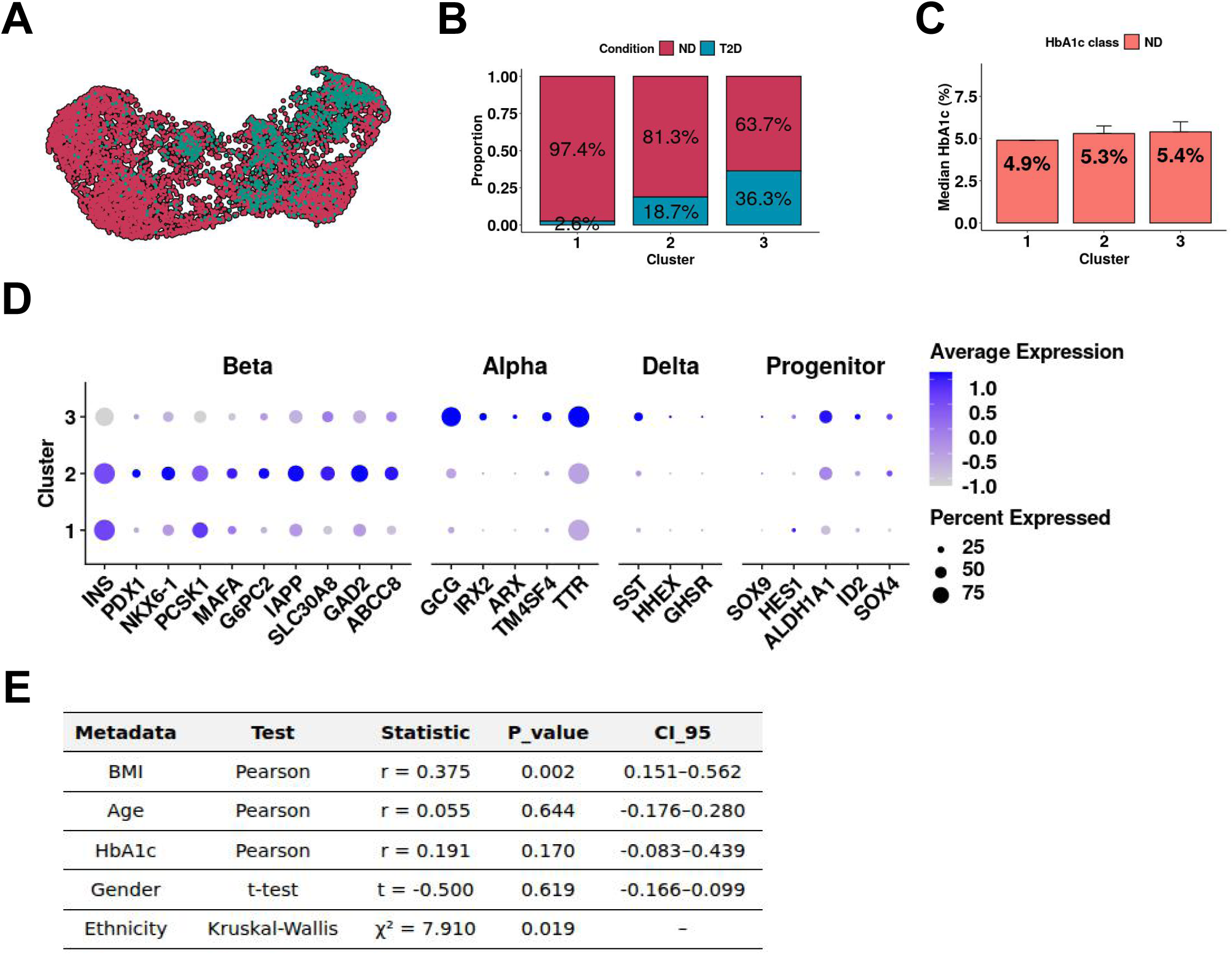
Molecular features and donor trait associations of the cluster C3 in Lineage3. (A) UMAP of Lineage3 beta-cells colored by disease condition (ND vs T2D). (B) Proportion of ND and T2D cells contributing to each cluster. (C) Barplot showing the median HbA1c per cluster with error bars representing MAD. (D) Dotplot showing scaled average expression and percent of cells expressing key beta (e.g., *INS, MAFA, PDX1*), alpha (e.g., *GCG, ARX*), delta (e.g., *SST, HHEX*), and progenitor-like (e.g., *SOX9, ALDH1A1*) markers across clusters. C3 exhibits co-expression of beta-cell and alpha/delta/progenitor-associated genes, consistent with poly-hormonal dedifferentiation. (E) Table summarizing associations between donor traits and C3 cluster occupancy. BMI shows a significant positive correlation (r = 0.375, P = 0.002), and ethnicity is also significantly associated (P = 0.019). No significant associations were observed for age, HbA1c, or gender.

**Figure S8:**
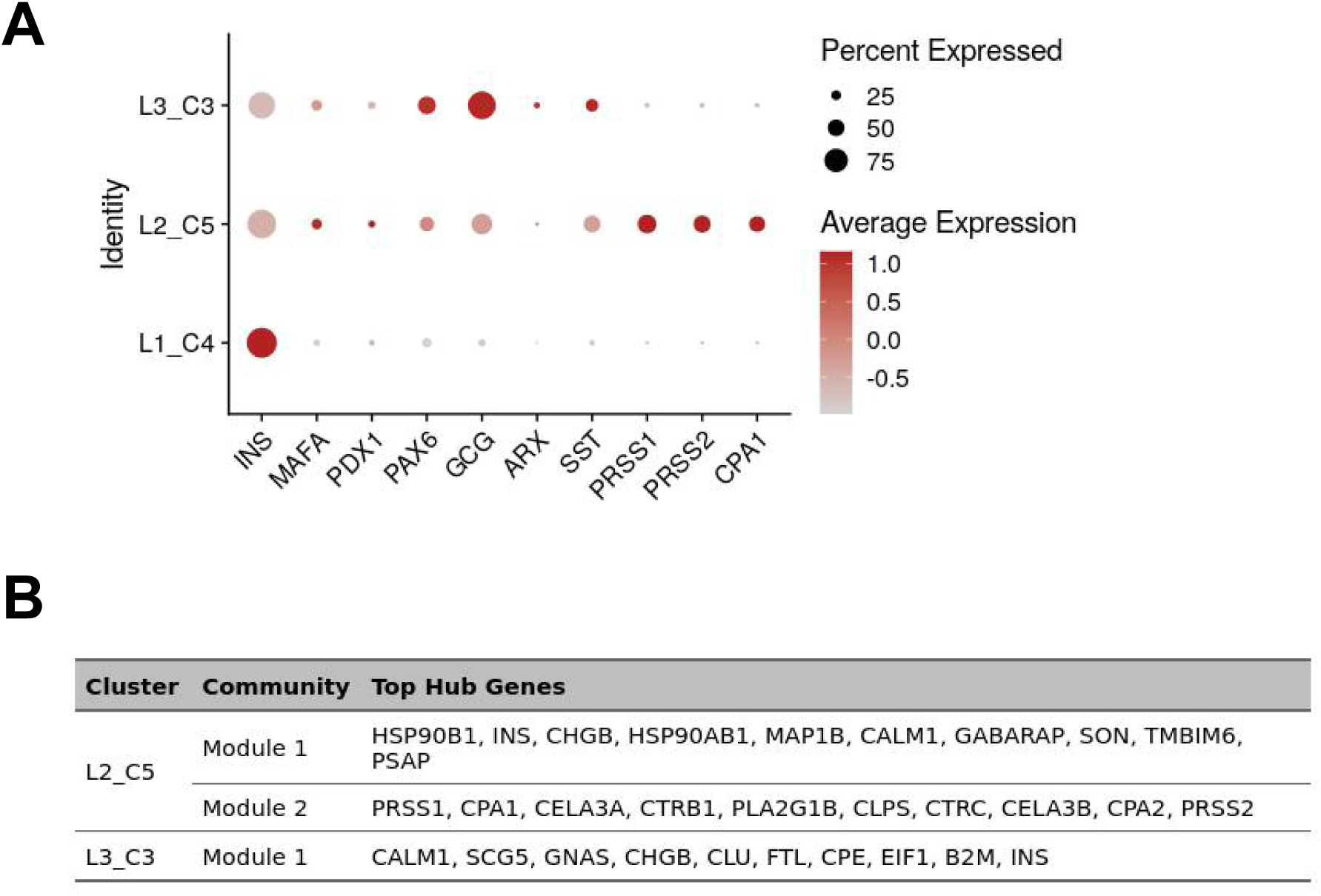
Extended analysis of divergent beta-cell identity-loss programs. (A) Dot plot of selected marker genes across L2_C5, L3_C3, and L1_C4 showing endocrine and exocrine genes expression. (B) Top hub genes of hypergraph modules identified by miEdgeR in L2_C5 and L3_C3.

## SUPPLEMENTARY TABLES

**TABLE S1. Summary of studies included in human pancreas scRNA-Seq atlas.**

**TABLE S2. Meta data of studies and individual datasets included in scRNA-Seq atlas.**

**TABLE S3. Summary of filtering settings used for pre-processing scRNA-Seq atlas datasets.**

**TABLE S4. ScRNA-Seq atlas differential gene expression analysis by cluster.**

**TABLE S5. Top 2000 most variable genes within each lineage.**

**TABLE S6. Lineage1_1 differential Start-vs-End test.**

**TABLE S7. Lineage1_2 differential Start-vs-End test.**

**TABLE S8. Beta-cell Lineage1 pseudobulk differential gene expression analysis of end trajectory clusters (cluster 3 versus cluster 4).**

**TABLE S9. Beta-cell Lineage2 pseudobulk differential gene expression analysis of end trajectory clusters (cluster 2 versus cluster 5).**

**TABLE S10. Lineage2_1 differential Start-vs-End test.**

**TABLE S11. Lineage2_2 differential Start-vs-End test.**

**TABLE S12. Lineage3 differential Start-vs-End test.**

**TABLE S13. Beta-cell Lineage3 pseudobulk differential gene expression analysis of end trajectory clusters (cluster 3 versus cluster 2).**

**TABLE S14. Module 1 gene overlap and cluster-specific components between L2_C5 and L3_C3.**

